# A qualitative study exploring researchers’ perspectives on authorship decision-making

**DOI:** 10.1101/615112

**Authors:** Lauren A. Maggio, Anthony R. Artino, Christopher J. Watling, Erik W. Driessen, Bridget C. O’Brien

**Affiliations:** Associate Professor of Medicine at Uniformed Services University of the Health Sciences in Bethesda, Maryland USA; Professor of Medicine at Uniformed Services University of the Health Sciences in Bethesda, Maryland USA; Professor of Clinical Neurological Sciences and Associate Dean for Postgraduate Medical Education at the Schulich School of Medicine and Dentistry, Western University, London, Canada; Professor of Medical Education, Faculty of Health, Medicine and Life Science, Maastricht University, the Netherlands; Associate Professor of Medicine at the University of California, San Francisco, San Francisco, California, USA

## Abstract

**Background:** Authorship has major implications for a researcher’s promotion and tenure, future funding, and career opportunities. Due in part to these high-stakes consequences, many journals require authors to meet formal authorship criteria, e.g. the International Committee of Medical Journal Editors (ICMJE) criteria for authorship. Yet on multiple surveys, researchers admit to violating these criteria, suggesting that authorship practices are a complex issue. Using qualitative methods, we aimed to unpack the complexities inherent in researchers’ conceptualizations of questionable authorship practices and to identify factors that make researchers vulnerable to engaging in such practices.

**Methods and Findings:** We conducted an interview study with a purposeful sample of 26 North American medical education researchers holding MD (n=17) and PhD (n=9) degrees and representing a range of career stages. We asked participants to respond to two vignettes – one portraying honorary authorship, the other describing an author order scenario – and then to describe related authorship experiences. Through thematic analysis, we found that participants, even when familiar with ICMJE criteria, conceptualized questionable authorship practices in various ways and articulated several ethical gray areas. We identified personal and situational factors, including hierarchy, resource dependence, institutional culture and gender, that contributed to participants’ vulnerability to and involvement in questionable authorship practices. Participants described negative instances of questionable authorship practices as well as situations in which these practices occurred for virtuous purposes. Participants rationalized that engagement in questionable authorship practices, while technically violating authorship criteria, could be reasonable when the practices seemed to benefit science and junior researchers. Participants described negative instances of questionable authorship practices as well as situations in which these practices occurred for virtuous purposes. Participants rationalized that engagement in questionable authorship practices, while technically violating authorship criteria, could be reasonable when the practices seemed to benefit science and junior researchers.

**Conclusion:** Authorship guidelines, such as the ICMJE criteria, portray authorship decisions as black and white, effectively sidestepping key dimensions that create ethical shades of gray. Our findings show that researchers generally recognize these shades of gray and in some cases acknowledge breaking or bending the rules themselves. Sometimes, their flexibility in applying rules of authorship is driven by benevolent aims that align with their own values or prevailing norms such as generosity and inclusivity. Other times, their participation in questionable authorship practices is framed not as a choice, but rather as a consequence of their vulnerability to individual or system factors beyond their control. Taken together, the findings reported here provide insights that may help researchers and institutions move beyond recognition of the challenges of authorship and contribute to the development of informed, evidence-based solutions for questionable authorship practices.

---

Authorship has high-stakes implications for researchers’ promotion and tenure, future funding, career opportunities, wellness, and sense of professional identity. Due in part to these consequences, questionable authorship practices and resultant authorship disputes are common [1-4]. Beyond the individual researcher, questionable authorship practices can have negative long-term effects for the scientific enterprise as a whole. These effects include distortion of the scientific record, dilution of authors’ true contributions, reinforcement of hierarchical abuse in academia, resentment within research teams, and negative impacts on patient care [5, 6]. To date, questionable authorship practices have primarily been studied using surveys and bibliometric analysis [7]. While valuable in estimating the frequency and types of questionable authorship practices in the field, these methods are limited in their capability to unpack the complex nature of authorship practices, thus providing little guidance on how to mitigate questionable actions and inactions.

Recognizing the gravity of authorship issues, journal editors have advocated authorship criteria to guide authors in planning projects and navigating authorship disputes [8]. The International Committee of Medical Journal Editors (ICMJE) have developed such authorship criteria and these have been widely adopted in biomedicine. According to ICMJE criteria, each author must: 1) Substantially contribute to the conception or design of the work; or the acquisition, analysis, or interpretation of data for the work; 2) Draft the work or revise it critically for important intellectual content; 3) Give final approval of the version to be published; *and* 4) Agree to be accountable for all aspects of the work in ensuring that questions related to the accuracy or integrity of any part of the work are appropriately investigated and resolved [9].

While a journal’s authorship criteria may be informative in articulating expected norms, the prevalence of questionable practices suggests they are insufficient to prevent authorship abuses. From studies that report evidence of questionable authorship practices [7, 10-12], it has become clear that authorship is a complex issue that requires a multi-faceted approach beyond simply knowing what constitutes authorship. The variety of questionable authorship practices reflect this notion. For example, when an individual has not satisfied the journal’s criteria for a given publication, his/her inclusion as an author is a questionable authorship practice, commonly referred to as honorary authorship. Honorary authorship may be given to an individual for a variety of reasons, such as the belief that adding a well-known author might enhance the work’s odds of being published, or as a reward for a collaborator who has contributed funding and/or resources to the research (so-called gift authorship) [2]. So, while the ultimate action – honorary authorship – might be the same, the rationale behind the action likely differs from one paper to another. In addition to honorary authorship, several other practices fit into the general category of questionable authorship practices, including, but not limited to, the inappropriate use of positional power to demand authorship or a particular location in the author by line, and the exclusion of authors who have significantly contributed (so-called ghost authorship)[13, 14].

Reacting to the variety of questionable authorship types, Moffatt recently proposed that the multiple ways in which researchers and their fields of study define and sanction authorship practices can contribute to conceptual confusion about authorship that negatively impacts ethical norms [15]. Yet, we know little about how researchers conceptualize questionable authorship practices, thereby making it difficult to offer the clarity needed to reduce confusion and negative impact. Thus, in this study we used interviews to begin to unpack the complexities inherent in researchers’ conceptualizations of questionable authorship practices and to further understand personal and situational factors that make individuals, and perhaps their research units and institutions, vulnerable to questionable authorship practices.

## Method

We conducted a qualitative interview study using thematic analysis guided by a constructivist approach. This study received an exempt determination from the Uniformed Services University of the Health Science’s Institutional Review Board (protocol #HU-MED-83-9684) and was approved by the Netherlands Association for Medical Education Ethics Review Board (dossier #1039). Per the regulations of these two bodies, access to the interview data is strictly controlled and limited to the core research team making it impossible for us to publicly deposit this data or make it available upon request. Moreover, the size and nature of our sample makes it possible that participants could be potentially identified despite anonymization of transcripts.

Recruitment, data collection, and analysis occurred between May 2018 and December 2018. We focused recruitment on the multidisciplinary field of medical education. Medical education researchers hail from a variety of academic backgrounds and research traditions (e.g., clinical medicine, psychology, biomedicine, education). We believe that the inclusion of a broad spectrum of researchers within a single field broadens the applicability of our findings. We purposively sampled participants who had published a multi-authored medical education research study between 2016 and 2017. To identify participants, we searched Web of Science (WOS) for articles published in *Academic Medicine* and *Medical Education*. We limited our initial search to these journals as they are the top two medical education journals based on impact factor. To broaden our sample, we also searched WOS for research articles in the 13 *JAMA* family journals using the key words “medical education” OR “health professions education” (see supplemental Appendix 1 for complete journal list). All included journals explicitly require that authors adhere to ICMJE guidelines.

Upon retrieval from all journals, we retained articles defined by WOS as research articles first authored by individuals based in the United States or Canada. We focused on North American authors due to what we perceived as similarities in university structures, such as promotion and tenure processes and hierarchy. Owing to the sensitive nature of the topic, we largely excluded authors from our own institutions although a single participant was from the same institution as one of the authors. From the included articles, we extracted the first author’s name and contact information. Our strategy yielded 119 potential participants, each of whom was invited by LAM a single time by email. Thirty-one potential participants responded to the invitation and were scheduled to be interviewed.

LAM conducted all interviews, between June and August 2018, via phone using a semi-structured interview guide. We piloted the interview guide with two researchers not in the study sample. Based on pilot feedback, we revised the guide accordingly prior to launching the study (See supplemental Appendix 2 for the final interview guide).

Interviews ranged in duration from 45 to 60 minutes. Prior to beginning the interview, informed consent was obtained from all participants. In the interview, LAM first asked participants to describe their reactions to two scenarios in which 1) their department chair added his/her name to the participant’s manuscript despite minimal contribution and 2) their position in the author order was shuffled from second to fourth author without notification. We opened the interviews with these two scenarios to provide participants with the same stimulus and starting point for considering questionable authorship practices. We selected these particular scenarios as they represent prevalent questionable authorship practices as reported in the literature [7, 10-12]. We believed the scenarios to present relatively clear and straightforward violations of authorship criteria. Following the discussion of these scenarios, participants were asked to describe any questionable authorship practices they had personally faced and how they handled those situations. Participants were also asked about their familiarity with the ICMJE criteria.

LAM conducted interviews in blocks of six participants. Upon completion of each interview block, recordings were transcribed, anonymized, and made available to the research team for discussion and preliminary analysis.

In alignment with qualitative methods, we stopped conducting interviews when the team agreed that we had reached data saturation. This occurred after 26 interviews. We defined data sufficiency as the point at which we could derive a clear and coherent understanding of key issues and could identify no additional nuances or insights to the issues. [16, 17]. Upon reaching data sufficiency, LAM thanked previously scheduled participants for their willingness to participate and cancelled their interview appointments.

We interviewed 26 (13 female) researchers based at 26 institutions in the United States (n=16) and Canada (n=10). Participants represented assistant (n=9), associate (n=8), and full (n=9) professors and held MD (n=17) and PhD (n=9) degrees. On average participants had published 56 journal articles (Range 1-301; SD=68.24).

To identify, analyze, and report patterns found in our transcripts, we utilized thematic analysis [18]. Analysis was primarily conducted by LAM and BCO beginning with line-by-line reading of transcripts to identify and define potential codes. In this close reading, which began after the first six participant interviews, we identified preliminary codes and working definitions related to factors described by participants. LAM and BCO coded all transcripts using Dedoose (Los Angeles). To complement LAM and BCO’s efforts, throughout the data collection, the other members of the research team actively reviewed transcripts, considered and discussed the resonance and fit of the identified codes, thereby helping to raise the level of analysis from categorizing to conceptualizing.

The first author assembled the research team based on members’ topical interests and the variety of roles each plays within the scientific community to inform the conceptualization of the study and data analysis. Our research team included five faculty members all holding PhD degrees. CJW is also a practicing physician in Canada. Combined, the team has engaged in authorship discussions for over 300 publications, served as editors and editorial board members, and held leadership responsibilities in graduate programs in schools of medicine. BCO and LAM are associate professors and ARA, CJW, and EWD are full professors. EWD is also a department chair.

## Results

Participants conceptualized questionable authorship practices in various ways and articulated several ethical “gray areas.” All participants described diverse personal experiences with and reactions to questionable authorship practices, including those presented in the two scenarios. Personal and situational factors contributed to participants’ perspectives on and involvement in questionable authorship practices. Some factors were sources of vulnerability; others served protective functions. Over time and the course of a career, some factors initially associated with vulnerability became protective factors. Below, we elaborate on researcher conceptualizations and factors contributing to or protecting against vulnerability, using representative quotations from a variety of participants.

### Conceptualizations of questionable authorship practices

The ICMJE criteria offer a way for authors to determine whether or not their contributions, and those of their colleagues, warrant authorship. Participants’ reactions to the presented scenarios, and the examples they related from their own experiences, revealed that even when they were familiar with the ICMJE criteria, as most were, their sense of what contributions warrant authorship varies. For example, some participants reacted to the scenario in which their chair added his/her name to the list of authors as a clear ethical violation: “To go against established understanding of rules of authorship is unprofessional… The most important ethical issue is that of professionalism and disrespect for the precepts on which we base our work as professionals. Also called lying.” (A)

Others considered additional criteria that could make their chair’s behavior less clear cut, providing a rationale that might make a decision to honor authorship less questionable or unethical. For example, several participants rationalized that if the chair had provided funding or had mentored the researcher, even if not in relation to the manuscript in question, then authorship might very well be warranted. In more than one case, participants admitted that they would actively seek out a potential justification to include the chair. For example, one participant said, “I would be looking for an answer to find a case in which I wouldn’t need to have that very uncomfortable conversation with the chair. I’d love to find some evidence of circumstances that would justify the inclusion.” (K)

As participants delved into the details of a variety of their own authorship situations, we found many were complex, nuanced, and difficult to evaluate based on the ICMJE criteria alone. In several situations, some participants described circumstances in which good intentions could underlie questionable authorship practices; that is, while the actions might technically violate the criteria for authorship, they might also be perceived to benefit the community and its researchers. For example, several participants rationalized engaging in questionable authorship practices to contribute to the greater good, especially when their actions benefitted junior researchers. When discussing the scenario on authorship order, one senior participant responded:

> I might also find out that this move was an attempt to help two junior authors in their careers who would really benefit. So that this was done not as a reflection of my contribution, but really to address another value, which I would agree with, which is helping a junior faculty. (K)

Another participant likened their department’s efforts to include junior researchers, even if they did not meet all ICMJE criteria, as a necessary form of “academic socialism” (C) to enable academic success for their junior colleagues, in light of their heavy clinical and teaching responsibilities and the challenges associated with promotion and tenure requirements.

Participants also indicated that the desire to act for the greater good – while not always strictly aligned with ICMJE criteria – supported the broader ethos of their scientific community.

> Intellectual communities are really feisty and individualistic and fight tooth and claw for dominance…We don’t want to be these people. We want to be caring and thoughtful. We want to bring people in and scaffold them and say ‘oh so this is your first publication and, you know, you have just reviewed the manuscript, but that’s okay you can be a first author this time.’ (Q)

In addition to casting questionable authorship practices as both vice (i.e., unethical) and virtue (i.e., benefiting the greater good), participants described other authorship “gray zones” (F) and pointed out that the ICMJE criteria are silent on many of these more nuanced points. These gray zones encompassed confusion around including authors whose level of contribution may have shifted over the course of a project, due, for example, to illness or maternity leave; participants who engaged in a single component of a project, such as conducting statistical analysis only; and authors who were unable to fully contribute due to lack of knowledge or skill in a given area. One participant’s example illustrates the challenges these gray zones might create for authorship decisions:

> I work with people in the international setting and their English is really bad and so their ability to contribute to the writing is practically non-existent. So they don’t contribute a whole lot to the writing…but it is only because they couldn’t, not because they didn’t contribute to the study in many other ways or deserve to be authors. So, yes there are gray areas, but in these cases it is really about capability. (F)

### Factors contributing to authors’ vulnerabilities

While participants identified nuance and gray areas in authorship practices, most also related experiences that they clearly considered to be problematic or harmful. In exploring these experiences, we identified situational and personal factors that contributed to participants’ vulnerability to questionable authorship practices. These factors included hierarchy, resource dependence, institutional culture, and gender. In some circumstances, participants consciously chose to engage in these practices based on a perception or recognition of their own vulnerability; in other circumstances, they felt victimized by others’ practices.

Hierarchy both contributed to perceived vulnerability and served as a way to rationalize questionable authorship practices. Several participants described hierarchy as a power differential between junior and senior faculty. Junior faculty generally associated hierarchy with diminished power and lack of voice. When reacting to the chair scenario, a junior researcher explained:

> In this situation you are telling someone in power that you don’t think their contributions warrant authorship. I think that’s a hard conversation up the hierarchical slope. At a minimum it’s going to be uncomfortable, but in a bad situation I could imagine real life repercussions in terms of losing opportunities if you challenge the authority figure. (O)

In response to the same scenario, one senior researcher noted: “The hierarchy thing puts a lot of people in a position of inability to speak up. I fear so much for junior people in this and perhaps in many institutions that have no voice because they will get destroyed.” (A)

Within the context of hierarchy, several participants noted that researchers’ inexperience and lack of familiarity with cultural norms could increase their vulnerability to questionable authorship decisions. For example, a junior researcher said:

> I can imagine being more bendable at an early stage. I would be driven by fear that I would be fired. I don’t think I would have any idea what my rights are and what the chairman’s rights are and are not. So I think I’d be confused about whether this was something that is typical or not. (J)

Resource dependence was also presented as a vulnerability. Participants discussed pressure to include individuals as authors who had provided access to resources, or helped to secure resources, despite not meeting ICMJE criteria. Specifically, some participants described including authors who were initially involved in attaining grant funding.

> I’ve carried 12 people on an authorship byline and two of them never made any contributions to the writing, but they had their names on the funding that funded the study. It was crazy - they never contributed anything. I had never seen them. I’m not even sure they know what I look like and I’ve given them several papers because they brought in the money. (Q)

Adding authors who provided access to research data or study populations necessary for study completion was also described. For example, one participant described an instance in which a program director demanded authorship on a survey study that included data from his residents, even though he contributed nothing to the conceptualization or writing of the manuscript. When describing a similar situation, a participant reported feeling like a “data hostage.”(C) Data access issues were frequently mentioned in relation to multi-institutional collaborations and when implementing research projects that were longitudinal, included multiple stakeholders, or cut across programs/departments.

> So I think it’s access to the data, but it is also keys to the kingdom that you need to make your work work…In order to operationalize the idea you need the buy in of a course director or someone that needs to do some nominal administrative thing and they are not really willing to do those nominal things unless there is the carrot of authorship. (C)

Participants also identified a high-pressure institutional culture as a situational factor related to (1) demanding messaging from leaders who articulated the need for faculty to publish, especially as first or last author; (2) poor role modeling of authorship behaviors; and (3) tacit institutional endorsement of questionable authorship practices. One participant noted:

> At a lot of institutions, I think there is such pressure for junior faculty to do well and to publish and to advance that it becomes oppressive. I think under that system of oppression people may end up doing things that they normally wouldn’t do or that they aren’t ethically comfortable with, but they need to do them out of necessity… I think what we are saying in a way is that this pressure is leading to a system or climate that leads to compromised ethical ways of being. (N)

Related to institutional culture, a participant shared a recent salary-related development at their university which they felt would negatively impact authorship practices.

> The administrators basically said we are now going to pay you based on a formula that looks at the number of your publications a year. So now instead of getting an envelope of funding which would guarantee me time out of clinic I have to worry about how many publications I get in a year. (D)

Gender was also perceived as a potential vulnerability. In several instances, participants said that being female influenced their ability and willingness to advocate for authorship, receive opportunities to earn authorship, and speak up against what they felt were injustices. For example, one female participant stated: “Gender discrimination against women in academia is alive and well, and I think a lot of junior women suffer the brunt of these abusive authorship practices because they are disempowered compared to the dominant men in the field” (Q). Several participants also described that they or their female colleagues had “no voice” (L) to speak out against honorary authorship practices as female researchers. Participants of both genders described this factor as a vulnerability for women.

### Protection from vulnerability

While we have focused on vulnerabilities, some participants also discussed protective elements that could help counterbalance these vulnerabilities. For example, several participants described components of their institutional processes and culture as protective. Such protective institutional factors included researcher training, resources to assist in authorship decisions (e.g., an ombudsperson), and strict enforcement of authorship standards.

> My institution has workshops…from an institutional standpoint this institution has tried to send messages about the importance of truly meeting guidelines of authorship and not just adding names to add names to papers. I think these workshops have contributed to a culture that values that type of integrity. (M)

Several physician participants mentioned their salary source, which is often linked to patient care and not research output, as protective and influential in their willingness to speak up about questionable authorship practices, or to not be concerned about them. Commenting on the scenario in which a department chair attempts to claim undue authorship, one physician participant noted:

> I’m an MD. Almost none of my income is controlled by my chair…So she could be mad at me, but unless she was so mad at me that she trash talked me to the dean it’s hard to imagine her having much control.” (AA)

We observed some examples of factors that seemed to shift over the course of a career, moving from a vulnerability to a source of protection; these included movement up the organizational hierarchy and greater experience. For instance, several senior participants reflected on personal experiences in which they felt that their lack of power as a junior researcher influenced their decision to engage in questionable authorship incidents – an effect that changed as they attained greater experience. One participant reflected on a personal experience from decades earlier:

> I did a research project on hemophilia and I worked with one guy closely and we did everything. Then in the last drafts of the paper there was some other guy’s name on it. I said, ‘who is this?’. Oh that is so and so and he runs the hemophilia lab. And you know being a PGY-2 at that time, I didn’t say anything…I think as you get older in your career you realize well no it’s not the way it should go. You now have a little bit more security and start to challenge. (D)

Additionally, senior researchers noted that their years of experience could be protective in that it increased their awareness of questionable authorship practices. One participant stated, when discussing the scenario on authorship order, “I’m just much more aware of these issues than someone who is diving into it brand new. So now I think about it right up front and address it with people when we start doing research”(E).

## Discussion

Despite over 1,000 journals requiring authors to adhere to ICMJE or similar criteria, our results underscore the idea that these criteria are insufficient for many of the complex and nuanced circumstances encountered in practice. Our results suggest that researchers’ conceptualizations of questionable authorship practices vary in intent, perceived consequences, and surrounding circumstances. Researchers’ experiences of such practices are influenced by multiple personal and situational factors such as hierarchy, resource dependence, experience, institutional culture, and gender. Some of these factors make researchers vulnerable to questionable authorship practices. In the sections that follow, we explore the implications of our findings, as well as some opportunities to address the complexities of authorship practices.

Most authorship guidelines (e.g., ICMJE criteria) portray authorship decisions as black and white, effectively sidestepping key dimensions that create ethical shades of gray. Our findings show that researchers generally recognize these shades of gray and in some cases acknowledge breaking or bending the rules themselves. Sometimes, their flexibility in applying rules of authorship is driven by benevolent aims that align with their own values or prevailing norms such as generosity and inclusivity. Other times, their participation in questionable authorship practices is framed not as a choice, but rather as a consequence of their vulnerability to individual or system factors beyond their control. While researchers recognize these latter breaches as potentially harmful to individuals, institutions, and the scholarly enterprise, they tend to characterize the former breaches as potentially helpful, supporting the progress of individual colleagues and nurturing the larger research community in the process. Thus, we see decision-making around authorship as far more complex than a simple application of guidelines. Rather, researchers navigate tensions between guidelines, their own values, and the factors that either protect them from or make them vulnerable to pressures from the often-hierarchical systems in which they work.

But when authorship decision-making is driven by individual circumstances and institutional culture, the resulting decisions may be idiosyncratic and inconsistent, threatening the integrity of the resulting scholarship. Our results do highlight gray areas where flexibility may be needed, such as circumstances where individual authors make very meaningful contributions to a piece of work but are unable to meet all of the authorship criteria. But they also point to authorship decision-making that may be motivated by a misguided sense of serving the greater good. Surely a junior faculty member will be better served by being mentored to undertake all the tasks that would earn first author status, for example, rather than being gifted with the status simply because their colleagues feel it will help their career.

We contend that researchers need more than guidelines; they require resources that assist in the situational interpretation of guidelines, mentorship in how authorship decisions can be made consistently and fairly, and education in the distinctly challenging communication skills needed to lead and participate in authorship discussions. They also require leadership from their more senior colleagues and a firm institutional commitment to integrity around authorship. And that commitment must not be undermined by a hidden curriculum that values productivity over authorship ethics.

This study should be considered in light of its limitations. While we purposively sampled researchers, there may be some degree of selection bias in our sample; that is, individuals who volunteered to be interviewed may be different in important ways from those who did not. That said, it is important to note that we collected a spectrum of responses from our participants, including those who reported direct engagement with such practices and those who had not experienced them. Another important limitation relates to participants’ willingness to share experiences with us. While all participants acknowledged the existence of questionable authorship practices, none admitted to acting as a “perpetrator” of clearly unethical authorship practices. Social desirability bias may in part explain this finding. Next, we only sampled medical education researchers, which limits the extent to which these findings might generalize to researchers in other scientific disciplines. Nonetheless, because medical education is a multi-disciplinarily field, our sample included researchers from a range of disciplinary backgrounds, and thus we believe the findings offer a broad perspective on the problem. Finally, we focused only on North American researchers; given the perceived influence institutional culture, the issues related to authorship in non-North American cultures might be distinct and offer a potential area of future research.

## Conclusion

While formal authorship standards like the ICMJE criteria provide guidelines for researchers, they appear inadequate to consistently mitigate questionable authorship practices. Our findings suggest that questionable authorship practices are complex, variably conceptualized by researchers, and situation-dependent, and thus somewhat resistant to a one-size-fits-all solution. We believe our findings provide insights that may help researchers and institutions move beyond recognition of the problem and contribute to the development of informed, evidence-based solutions for questionable authorship practices.

## Funding/Support

No funding was received for this work.

### Acknowledgements

The authors wish to thank all of the participants for sharing their insights and stories.

## Disclaimer

The views expressed in this article are those of the authors and do not necessarily reflect the official policy or position of the Uniformed Services University of the Health Sciences, the Department of Defense, or the U.S. Government.

## Supplemental Appendix 1: Complete list of journals searched to identify first authors of research studies

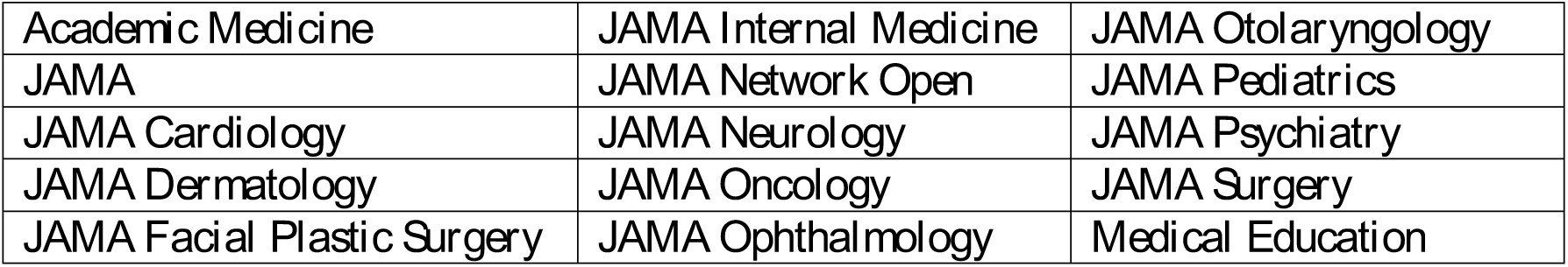

|  |  |  |
| --- | --- | --- |
| *Academic Medicine* | *JAMA Internal Medicine* | *JAMA Otolaryngology* |
| *JAMA* | *JAMA Network Open* | *JAMA Pediatrics* |
| *JAMA Cardiology* | *JAMA Neurology* | *JAMA Psychiatry* |
| *JAMA Dermatology* | *JAMA Oncology* | *JAMA Surgery* |
| *JAMA Facial Plastic Surgery* | *JAMA Ophthalmology* | *Medical Education* |

## Supplemental Appendix 2

Semi-structured interview guide

To get started:

I’m going to read you a short scenario. Please think through it and describe how you would react if you were in this situation.

### Scenario 1

- You are the lead author of a study. You have discussed the study with your department chair/head several times and have kept her updated on your progress. Just prior to submitting the article for peer review, your chair offers to provide feedback. You send her your final draft and it is returned the next day. You see a few grammatical suggestions and notice that your chair has added her name as the last author.

How would you react in this situation?

What thoughts and/or questions came to you as you listened to the scenario?

Follow-ups

- Is there an ethical problem here and, if so, what is it (in your estimation)?
- How might relationships play into this situation?
- What, if any, elements of power might come into play in this situation?
- What potential cultural elements (e.g., HPE authorship conventions, local institutional practices) might impact your approach to this situation?
- What potential personal factors (e.g., seniority, gender, educational background, relationship to the individual/your author team, promotion and tenure) might impact your approach to this situation?
- What resources, including colleagues/mentors/ombudsmen, might you have available to you to facilitate your handling of this situation?
- In this scenario your chair listed herself as last author. In what ways if any did that influence your thinking?

#### Scenario 1: Personal experience

Have you ever found yourself in a similar situation? If yes, please describe the situation and how you handled it.

Follow-ups:

- What thoughts, emotions, and/or other considerations did you experience?
- IN this experience did you provide your co-authors with feedback?
- How might relationships play into this situation?
- What, if any, elements of power might come into play in this situation?
- What potential cultural elements (e.g., HPE authorship conventions, local institutional practices) impacted your approach to this situation?
- What potential personal factors (e.g., seniority, educational background, relationship to the individual/your author team) impacted your approach to this situation?
- What elements of power were at play in this situation?
- What resources, including colleagues/mentors/ombudsmen, did you have available to facilitate your handling of this situation?
- In what ways might you handle this situation differently now and why?

In the first scenario, we discussed some of the challenges related to academic publishing and authorship. In what ways, if any, do you feel that this practice can be damaging?

- For the field of HPE?
- For individual authors?
- For you personally?

### Scenario 2

Okay, I’m going to read you one more short scenario. Just like last time, please think through it and describe how you would react if you were in this situation.

- For the last six months, you have been working closely with a team of senior researchers to publish a literature review. Your work has included reviewing a portion of the literature, assisting with the coding, and writing sections of the review paper. At the beginning of the project you were promised that you would be the second author and feel that your efforts warrant such a position. When it comes time for submission, the first author circulates the manuscript and you notice that you have been bumped to fourth author.

How would you react in this situation?

What thoughts and/or questions came to you as you listened to the scenario?

Potential probes (if needed):

- Is there an ethical problem here and, if so, what is it (in your estimation)?
- What potential cultural elements (e.g., HPE authorship conventions, local institutional practices) might impact your approach to this situation?
- How might relationships play into this situation?
- What, if any, elements of power might come into play in this situation?
- What potential personal factors (e.g., seniority, educational background, gender, relationship to the individual/your author team) might impact your approach to this situation?
- What resources, including colleagues/mentors/ombudsmen, might you have available to you to facilitate your handling of this situation?

#### Scenario 2: Personal experience

Have you ever found yourself in a similar situation? If yes, please describe the situation and how you handled it.

Potential probes (if needed):

- What thoughts, emotions, and/or other considerations did you experience?
- What potential cultural elements (e.g., HPE authorship conventions, local institutional practices) impacted your approach to this situation?
- What potential personal factors (e.g., seniority, educational background, relationship to the individual/your author team) impacted your approach to this situation?
- What elements of power were at play in this situation?
- What resources, including colleagues/mentors/ombudsmen did you have available to facilitate your handling of this situation?
- In what ways might you handle this situation differently now and why?

In this scenario, we have discussed issues related to authorship order. In what ways, if any, do you feel that current practices related to author order can be damaging?

- What about the field of HPE?
- What about for authors?
- What about to you personally?

### General Questions

Thank you for walking through those scenarios with me. I have a few more questions.

Recently you were the lead author on a published a paper in Academic Medicine/Medical Education/JAMA.

For that article, did you have an explicit conversation about authorship for that paper? If yes, (If no, can you describe an experience in which you did have an explicit authorship conversation?)

- How did you initiate the conversation about authorship?
- What topics did you cover?
- At what point did you initiate this discussion?
- For each author were there any particular characteristics (e.g. methodological skills, access to data, reputation) that led you to invite them to the author team?
- How did you take into consideration the composition of the research team (e.g., a team of mostly senior scholars vs. a mixed team of senior and junior scholars)?
- How did you approach the topic of author order?
- At what points, if any, did you revisit the topic of authorship with your research team?
- Was there anything that surprised you about this authorship experience or something in particular that you would like share about it?
- In your experience, would you describe the authorship experience with this article as standard for your field? (or your research? Or the groups you work with? Or your institution/dept?)

More generally, can you describe:

- Who and/or what influenced your approach to discussing authorship?
  - Did you receive any formal training or mentoring?
- Can you describe a situation in which you rejected or turned down authorship that was offered to you?
- In some situations, projects recruit the assistance of a research assistant or a statistician. Have you had experience integration these colleagues onto a project and how did it go? How did you recognize them for their efforts?
- Can you describe a time when you felt someone who was initially a part of an authorship team no longer warranted authorship and how you handled the situation?
- Have you published outside of HPE? If yes, how did the experience contrast with experiences publishing in HPE?
- Researchers have described authorship as academic capitol. What are some ways that you think academic researchers use authorship as capitol?
- What responsibilities come with being an author?

In some scholarly circles, authorship has been defined by the four following criteria that state an author must:

1. Substantially contributes to the conception or design of the work; or the acquisition, analysis, or interpretation of data for the work; AND
2. Drafts the work or revises it critically for important intellectual content; AND
3. Gives final approval of the version to be published; AND
4. Agrees to be accountable for all aspects of the work in ensuring that questions related to the accuracy or integrity of any part of the work are appropriately investigated and resolved.

- Are you familiar with these criteria?
- To what degree do you think these criteria are applied in HPE research? What factors do you feel might influence how these criteria are applied?
- Beyond these criteria, are there any authorship practices (formal or informal) at your institution? How did you learn about these practices?
- Think about the last paper on which you were an author. Do you feel that you met all four criteria? What about your co-authors?
- Do you feel that there should be exceptions to these criteria? If yes, please describe an example.

In HPE, what can be done to help researchers navigate authorship decisions?

Okay then, I would like to close with a few demographic questions:

1. On a scale of 1-10 with 1 being a junior researcher, 5 a mid-career researcher and 10 a senior researcher, how would you describe your current level as an HPE researcher?
2. What is your academic rank?
3. Is tenure available at your institution? Have you attained tenure?
4. Please describe your educational background and include any specific training you have received in HPE.
5. In the Academic Medicine/JAMA/Medical Education paper, what is your primary identification?
6. How many years have you been doing HPE research?
7. What percentage of your average work week is dedicated to HPE research?
8. Approximately how many journal articles have you published?
  a. Of those how many were on HPE-related topics?
9. Would you characterize yourself as a qualitative, quantitative or mixed methods researcher?

Do you have any questions or additional comments that you would like to make about anything that we discussed today?

Those are all the questions that I have for you. Thank you for your time.

